# Identification of two novel polygalacturonase-inhibiting proteins (PGIPs) and their genomic reorganization in chickpea (*Cicer arietinum*)

**DOI:** 10.1101/2023.03.26.534275

**Authors:** Vishnutej Ellur, Wei Wei, Rishikesh Ghogare, Shyam Solanki, George Vandemark, Robert Brueggeman, Weidong Chen

**Affiliations:** Molecular Plant Science, Washington State University, Pullman, WA, USA; Department of Plant Pathology, Washington State University, Pullman, WA, USA; Department of Horticulture, Washington State University, Pullman, WA, USA; Department of Agronomy, Horticulture & Plant Science, South Dakota State University, Brookings, SD, USA; Grain Legume Genetics Physiology Research, USDA ARS, Pullman, WA, USA; Department of Crop & Soil Science, Washington State University, Pullman, WA, USA

**Keywords:** Polygalacturonase inhibitory proteins (PGIPs), gene family, defense-related gene, biotic stress response, leucine-rich repeats (LRRs), promoter analysis, constitutive gene expression, subcellular localization

## Abstract

Polygalacturonase inhibiting proteins (PGIPs) are cell wall-anchored proteins that inhibit pathogen polygalacturonases (PGs). PGIPs, like other resistance genes, contain extracytoplasmic leucine-rich repeats (eLRRs), which are required for pathogen PG recognition. The importance of these PGIPs in plant defense has been well documented. This study focuses on chickpea (*Cicer arietinum*) PGIPs (CaPGIPs) owing to limited information available on this important crop. In this study, we identified and characterized two novel *Capgips* (*Capgip3* and *Capgip4*) in addition to the previously reported *Capgip1* and *Capgip2*. Our analysis showed that CaPGIP1, CaPGIP3, and CaPGIP4 proteins contain N-terminal signal peptides, ten LRRs, theoretical molecular mass, and isoelectric points like other legume PGIPs. However, the previously reported CaPGIP2 cannot be classified as a true PGIP since it lacked a signal peptide, more than half of the LRRs, and other characteristics of a typical PGIP. Phylogenetic analysis and multiple sequence alignment revealed that the *Capgip* amino acid sequences are similar to the other reported legumes. Several cis-acting elements that are typical of pathogen response, tissue-specific activity, hormone response, and abiotic stress-related are present in the promoters of *Capgip1, Capgip3*, and *Capgip4*. Localization experiments showed that *Capgip1, Capgip3*, and *Capgip4* are located in the cell wall or membrane, whereas *Capgip2* is found in the endoplasmic reticulum. *Capgip* transcript levels analyzed at normal conditions show constitutive tissue specific expression and heterozygous expression patterns analogous to other defense related gene families. All these findings suggest that CaPGIPs could have the potential to combat chickpea pathogens.

## 1 Introduction

Plants deploy a variety of barriers to withstand numerous pathogenic stresses, one of which is the cell wall, a physical barrier that serves as the first line of defense. Pathogens produce enzymes known as cell wall-degrading enzymes (CWDEs) to overcome this plant barrier (Kubicek et al., 2014). Pectin-degrading enzymes polygalacturonases are among the most important CWDEs. Pectin is a key component of plant cell walls that determines the integrity and rigidity of plant tissue (Voragen et al., 2009), and degrading pectin enables quick access to the components within the cell. Pectin is majorly comprised of D-galacturonic acid. Polygalacturonases (PGs) are fungal enzymes that degrade pectin by breaking down glycosidic bonds between D-galacturonic acid residues and aid in tissue maceration. Polygalacturonases (PG) are secreted at the early stages of infection (De Lorenzo et al., 2001). In defense, plants use polygalacturonase inhibiting proteins (PGIPs) to impede PGs’ pectin-depolymerizing activity. Plant PGIPs are located on the cell wall and their potential to suppress PG activity is correlated with plant disease resistance (Ge et al., 2019).

PGIPs are highly conserved proteins (Di Matteo et al., 2003). Most PGIPs are generally intronless, except a few that include a short intron (Kalunke et al., 2015). PGIPs, like many resistance gene products, contain extracytoplasmic type leucine-rich repeats (eLRRs) (Di Matteo et al., 2003, and Kalunke et al., 2014). PGIPs are composed of 10 incomplete LRRs of approximately 24 residues each, which are arranged into two β-sheets. β1 occupies the inner concave side of the molecules, while β2 occupies the outer convex side. These repeats form β-sheet/β-turn/α-helix containing LRR motifs. Motifs that occupy the β1 inner concave side are critical for interaction with PGs. Like other members of the LRR superfamily, PGIPs contain immune receptors required for plant innate immunity. PGIPs have been reported in every characterized plant species so far (Kalunke et al., 2015) and can thus be used as defense molecules in disease resistance to fungal pathogens (Hamera et al., 2014, and Zhou et al., 2017).

Albersheim and Anderson were the first to report *pgip* activity in 1971. The first *pgip* gene, however, was isolated in French beans twenty years later. Several PGIPs have been identified in several crops based on sequence identity since 1971. *Pgip* genes do not undergo large expansion and may exist as single genes (Di Giovanni et al., 2008) or clustered into small gene families (Ferrari et al., 2003). In legumes, *pgips* have been characterized in *Glycine max, Medicago sativa, Medicago truncatula, Phaseolus acutifolius, Phaseolus coccineus, Phaseolus lunatus, Phaseolus vulgaris, Pisum sativum*, and *Vigna radiata*. However, only *M. truncatula, V. radiata, P. vulgaris, and G. max’s* genome have more than one *pgip*. Pathogen PG inhibition by PGIPs is well established. The majority of the identified legume PGIPs inhibited fungal PGs, such as *G. max’s* GmPGIP7 (D’Ovidio et al., 2006, Frati et al., 2006, and Kalunke et al., 2014), *M. truncatula’s* MtPGIP1, MtPGIP2 (Song and Nam, 2005), *P. vulgaris’s* PvPGIP1, PvPGIP 2, PvPGIP3, PvPGIP 4 (Desiderio et al., 1997, D’Ovidio et al., 2006, and Frati et al., 2006), *P. acutifolius’*s PaPGIP2, *P. coccineus’*s PcPGIP2, and *P. lunatus’s* PlPGIP2 (Farina et al., 2009). However, PGIPs from *V. radiata*, VrPGIP1, and VrPGIP2 have shown to inhibit insect PGs (Kaewwongwal et al., 2017) and *P. sativum* PsPGIP inhibited nematode PGs (Veronico et al., 2010).

Some *pgips* are constitutively expressed, while others respond to external cues. Pathogens such as fungi, oomycetes, insects, and nematodes are known to induce *pgip* expression, as are phytohormones such as abscisic acid (ABA), indole-3-acetic acid (IAA), salicylic acid (SA), and jasmonic acid (JA) (Ferrari et al., 2003, Hou et al., 2015, and Hwang et al., 2010). *Pgip* expression is also triggered by wounding and oligogalacturonic acid treatments (Di Matteo et al., 2006, and Ferrari et al., 2003). Although *pgip* genes/gene families are constitutively expressed, their expression is tissue-specific and developmentally regulated (Li and Smigocki, 2016), studies conducted with basal transcript levels of *B. napus pgips* (Hegedus, et al., 2008), *P. vulgaris pgips* (Kalunke et al., 2011), *and C. papaya pgips* (Broetto et al., 2015) indicate *pgips* are expressed in normal conditions when plants are not stressed.

Currently eighteen PGIPs have been characterized from nine legume species, but major legumes such as chickpeas, peanuts, and lentil PGIPs remain uncharacterized. This study focuses on PGIPs of chickpeas because chickpeas are the world’s second most widely produced and consumed leguminous crop. Chickpeas have a high protein content (up to 40% protein by weight), are an excellent source of essential vitamins such as riboflavin, niacin, thiamin, folate, and the vitamin A precursor β-carotene, and have other potential health benefits such as lowering cardiovascular, diabetic, and cancer risks (Jukanti et al., 2012; Merga and Haji, 2019; Sharma et al., 2013). Two chickpea PGIPs (CaPGIP1 and CaPGIP2) were previously reported based on sequence searches of the chickpea genome. The goal of this study is to investigate and characterize PGIPs in chickpeas to gain a better understanding of their structural features, functional domains, regulatory elements, and genomic organization. *Capgip* genes were cloned, and their sequence features were evaluated in this study. The basal expression of all *Capgips* was explored. Our findings revealed that *Capgips*, like other legume *pgips*, had similar characteristics and can play an essential role in plant resistance against fungal diseases.

## 2 Materials and Methods

### 2.1 Sequence acquisition, phylogeny, and bioinformatics analysis

To identify PGIP homologs in the chickpea genome, a homology search was performed against the NCBI database using the amino acid sequences of previously known legume PGIPs. SignalP 5.0 was used to identify the presence of signal peptides in the candidate genes identified by the NCBI homology search (Armenteros et al., 2019). The molecular weight and isoelectric point (pI) were determined using ExPASy Server (Gasteiger et al., 2005). NetOGlyc version 4.0 server was used to analyze the putative N-linked glycosylation sites (Steentoft et al., 2013). The Swiss-Model server was used to build homology-based 3D models of *Capgips* (Waterhouse et al., 2018). Protein sequences of previously characterized legumes were aligned using Clustal W. Jalview was used for multiple sequence alignment with a conservation index of 50% (Waterhouse et al., 2009). A phylogenetic tree was generated using MEGA X (Kumar et al., 2018) with the neighbor-joining phylogenetic statistical method, Poisson model and other settings retained at default. The tree was bootstrapped 1000 times for robustness and *Cucumis sativus* PGIPs (CsPGIPs) were used as outgroup. MEGA X generated trees were visualized using the iTOL version 6.1.1 online tool (Letunic and Bork., 2021). The 1,500 bp upstream sequence for all Capgip sequences was analyzed for the putative cis-acting regulatory DNA elements using New PLACE (Higo et al., 1999).

### 2.2 Cloning and sequencing

The total RNA was extracted from the leaves of chickpea cultivar Dwelley using the RNeasy Plant Mini Kit (Qiagen, Hilden, Germany) from 100 mg samples in accordance with the manufacturer’s protocol. First strand cDNA synthesis and genomic DNA elimination were performed simultaneously using 5X All-In-One RT MasterMix, containing AccurT Genomic DNA Removal (Applied Biological Materials Inc, Richmond, Canada). RNA extraction, genomic DNA removal, and cDNA synthesis were performed on the same day to preserve sample RNA integrity. cDNA samples were stored at −80°C until use. Full-length ORFs were amplified with Phusion^®^ High-Fidelity DNA Polymerase (NEB, Ipswich, MA, USA) using gene-specific primer pairs (Supplementary Table 2) using the following protocol: initial denaturation at 98 °C for 30 seconds and 35 cycles of 98 °C for 10 seconds, 60 °C and 72 °C for 30 seconds each and followed by a final elongation at 72 °C for 8 minutes. Amplified PCR products with appropriate expected sizes were purified with the Monarch^®^ DNA Gel Extraction Kit (NEB). Purified PCR products were cloned into the pMiniT 2.0 vector (NEB) and transformed into DH10B high-efficiency *E. coli* competent cells (NEB) by electroporation. The plasmids were recovered from *E. coli* using PureYield™ Plasmid Miniprep (Promega, Madison, WI, USA), verified using Sanger sequencing (Laboratory of Biotechnology & Bioanalysis, Pullman, WA, USA), and compared to the GenBank sequences of *Capgip1* (XM_004504675), *Capgip3* (XM_004493500), and *Capgip4* (XM_012713804).

### 2.3 Subcellular localization

DeepLoc-1.0 (Almagro Armenteros et al., 2017) was used to predict subcellular localization based on *Capgip’s* protein sequences. The complete coding sequences (CDS) of *Capgip1, Capgip2, Capgip3*, and *Capgip4* were cloned into pEarleyGate 103 using the gateway cloning approach to determine their subcellular localization. After sequencing validation, these gateway plant expression vectors were transformed into *A. tumefaciens* strain EHA105. Using a blunt syringe, transformed EHA105 cultures harboring *Capgip-mGFP* plasmids were infiltrated into 4-week-old *N. benthamiana* leaves. A laser scanning confocal microscope (Leica SP-8) was used to examine and capture the fluorescence emitted by fusion proteins 72 hours after infiltration. GFP fluorescence was excited at 488 nm.

### 2.4 Plant materials

Chickpea (*Cicer arietinum*) cultivar Dwelley was grown in greenhouse conditions. Plants maintained in the greenhouse at 22 ± 2°C. Leaf, stem, root, flower, pod, and seed tissues were collected at different chickpea growth stages, which are mentioned in Table 3. Tissue samples (100 mg) were collected in three biological replicates and were immediately snap-frozen in liquid nitrogen and stored at −80°C.

**TABLE 1:** NCBI accession number, full-length cDNA size (bp), ORF size (bp), predicted protein size (aa), predicted signal peptide size (aa), theoretical molecular mass (kDa), and pI for all four *Capgips*.

| Name | NCBI<br>accession | Gene symbol | cDNA<br>(bp) | ORF<br>(bp) | Protein<br>(aa) | Signal<br>peptide<br>(aa) | Molecular<br>mass<br>(kDa) | Isoelectric<br>point<br>(pI) |
| --- | --- | --- | --- | --- | --- | --- | --- | --- |
| *Capgip1* | XM_004504675 | LOC101505245 | 1365 | 1041 | 347 | 36 | 37.65 | 9.05 |
| *Capgip2* | XM_027335237 | LOC101504619 | 523 | 492 | 164 | NA | 18.18 | 5.64 |
| *Capgip3* | XM_004493500 | LOC101499240 | 1313 | 1005 | 335 | 22 | 37.61 | 8.44 |
| *Capgip4* | XM_012713804 | LOC101499557 | 1273 | 987 | 329 | 20 | 36.19 | 6.64 |

**TABLE 2:** Putative hormonal and pathogenesis related *cis*-acting regulatory elements identified in the promoter regions of *Capgips*.

| *Cis* - element | Position |  |  | Signal Sequence | Function | References |
| --- | --- | --- | --- | --- | --- | --- |
|  | *Capgip1* | *Capgip3* | *Capgip4* |  |  |  |
| ABRELATERD1 | - | 287(-),288(+),1415(-) | 561(-),1132(-),1167(+) | ACGTG | Abscisic acid response | Nakashima *et al.*, 2006 |
| ABRERATCAL | - | 286(-),287(+),1414(-) | 560(-),1131(-),1166(+) | MACGYGB | Abscisic acid response | Kaplan *et al.*, 2006 |
| ACGTABRE | - | - | 559(-) | ACGTGKC | Abscisic acid response | Hattori *et al.*, 2002 |
| ARFAT | - | - | 857(+) | TGTCTC | Abscisic acid response | Nag *et al.*, 2005 |
| ARR1AT | 320(-), 600(+),622(+),<br>637(-),969(-),990(-),<br>1008(-),1023(-),1054(-),<br>1223(-) | 151(-),1293(-),1410(-),<br>1435(+),1488(-) | 98(-),113(+),190(-),245(-),<br>355(+),371(-),872(+),896(-),<br>978(+),1106(+),1193(-),<br>1196(+),1275(-), 1360(+),<br>1446(-) | NGATT | Bacterial response | Ross *et al.*, 2004 |
| BIHD1OS | 198(-),1337(-) | 483(+),661(+),809(-),<br>1044(-) | 1300(-),1369(+) | TGTCA | Pathogen response | Luo *et al.*, 2005 |
| GADOWNAT | - | - | 559(-) | ACGTGTC | Abscisic acid response | Nakashima *et al.*, 2006 |
| GAREAT | 123(+),144(+) | - | 166(+),741(-) | TAACAAR | Gibberellic acid response | Ogawa *et al.*, 2003 |
| GT1CONSENSUS | 209(+),236(+),283(+),3<br>56(+),402(-),422(-),<br>430(-),714(+),754(-),<br>843(+),1091(-),1092(-),<br>1130(-),1146(-),<br>1203(+),1369(-) | 26(+),33(+),54(-),<br>73(+),87(+),391(-),<br>429(+),477(+),575(-),<br>612(+),668(-),712(+),<br>1102(-),1116(-),1136(-),<br>1310(-),1332(+),1465(-) | 53(+),66(-),86(-),106(-),<br>279(+),314(+),330(-),787(-),<br>799(-),907(+),908(+),<br>1021(-),1022(-),1031(-),<br>1046(-),1456(-),1470(-) | GRWAAW | Salicylic acid response | Buchel *et al.*, 1999 |
| GT1GMSCAM4 | 209(+),422(-),1091(-),<br>1146(-),1203(+),1369(-) | 26 (+),87(+),391(-),<br>612(+),1136(-),<br>1310(-),1332(+) | 53(+),314(+),<br>799(-),1470(-) | GAAAAA | Pathogen response | Park *et al.*, 2004 |
| SEBFCONSSTPR<br>10A | - | 660(+),783(-) | 856(+) | YTGTCWC | Pathogen response | Boyle & Brisson, 2001 |
| T/GBOXATPIN2 | - | 1415(-) | 561(-),1132(-),1166(+) | AACGTG | Jasmonate response | Boter *et al.*, 2004 |
| WBOXNTERF3 | - | - | 412(-),780 (-),1060(-),1119<br>(+),1293 (+),1307 (+) | TGACY | Wound response | Nishiuchi *et al.*, 2004 |

**Table 3:**
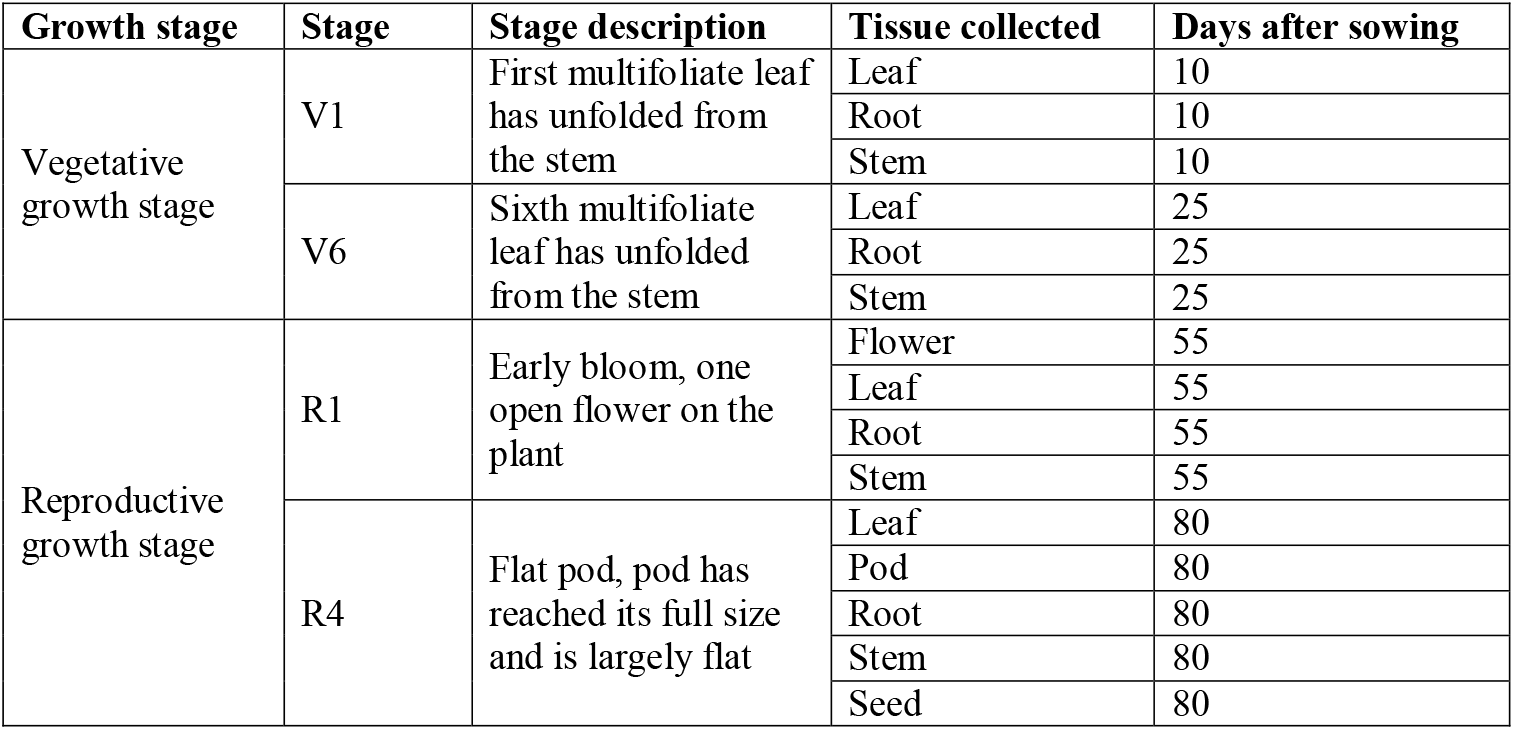
Chickpea tissues collected during different chickpea growth stages for absolute gene expression quantification.

### 2.5 RNA isolation, cDNA conversion and quantitative real-time-PCR

To determine the absolute expression of *Capgips* in normal conditions, total RNA was extracted from various chickpea tissues from the cultivar Dwelley at different growth stages (Table 3). RNA was extracted from 100 mg samples using the RNeasy Plant Mini Kit (Qiagen) according to the manufacturer’s protocol. First-strand cDNA synthesis and genomic DNA elimination were performed simultaneously using 5X All-In-One RT MasterMix, containing AccurT Genomic DNA Removal (Applied Biological Materials, Inc). To preserve sample integrity, RNA extraction, genomic DNA removal, and cDNA synthesis were performed on the same day. Samples were stored at −80°C until use. Using Primer3Plus (Untergasser et al., 2007) at the default parameters, quantitative PCR primers (Supplementary Table 1) were generated based on the sequences of *Capgip1, Capgip3*, and *Capgip4* and the reference gene 18SrRNA and 25SrRNA. The NCBI Primer-BLAST (ncbi.nlm.nih.gov/tools/primer-blast/) program was used to ensure that primers are unique specifically to the corresponding gene. Standard curves generated by serial dilution of cDNA for 18SrRNA, 25SrRNA, Capgip1, Capgip3, and Capgip4 were used to evaluate primer efficiency (Supplementary figure 1). Transcript levels of chickpea *pgips (Capgip1, Capgip3*, and *Capgip4*) in chickpea at normal conditions were evaluated following the Minimum Information for Publication of Quantitative Real-Time PCR Experiments (MIQE) guidelines (Bustin et al., 2009). Each RT-qPCR reaction consisted of 1x SsoAdvancedTM Universal Inhibitor-Tolerant SYBR^®^ Green Supermix (Bio-Rad, Hercules, CA, USA), 2.5 μM of each gene-specific primer, and 100 ng of cDNA in a final reaction volume of 10 μL. No template control, no amplification control, and negative reverse transcription controls were included for each primer pair, and all reactions were performed with three separate biological replicates in technical triplicates. qPCR was carried out in the CFX96TM Real-Time PCR Detection System using a two-step amplification and melt curve method with the following protocol: 95 °C for 3 minutes, followed by 40 cycles of 95 °C for 10 seconds; 60 °C for 30 seconds; and 72 °C for 30 seconds. Melt curve readings were taken from 65.0°C to 95.0°C with an increment of 0.5°C every 5 seconds. The absolute gene expression assays were performed by constructing standard curves of the corresponding cloned coding region of *Capgips* (Wong and Medrano, 2005). The expression Ct values of *Capgips* were normalized against the expression Ct values of reference genes 18SrRNA and 25SrRNA. Quantification was done using the relative standard curve method (Supplementary figure 1) (Pfaffl, 2001). The *Capgip* expression values are given as the mean of the normalized expression values of *Capgips* normalized against reference genes 18SrRNA and 25SrRNA. Obtained *Capgip* data is shown as gene copy number/microgram of RNA (Forlenza et al., 2012).

## 3 Results

### 3.1 Insilco characterization of *Capgips*

*Capgip1* (LOC101505245) and *Capgip2* (LOC101504619) are two previously reported chickpea *pgips* (Kalunke et al., 2014, and Kalunke et al., 2015). They occupy a 30,150 bp region on chromosome 6. *Capgip1* has a single exon with no intron as seen in several legume *pgips*. While *Capgip2* has two exons separated by a 9,825 bp intron. A homology search against the NCBI database using the amino acid sequences of known legume *pgips* revealed the presence of additional five candidate *pgip* sequences (Supplementary Table 1). Only two of them, LOC101499240 and LOC101499557, were suitable for designation as prospective *pgips* since they were the appropriate size, had a signal peptide, and had ten LRR sequences. They were named *Capgip3* and *Capgip4* respectively. *Capgip3* and *Capgip4* are composed of a single exon with no introns and span a 15,582 bp region on chromosome 3. In addition to the previously known locus of PGIP genes (Kalunke et al., 2014, and Kalunke et al., 2015), our analysis identifies a new locus. The occurrence of two *pgip* loci in chickpeas, chromosome 3 and chromosome 6, necessitates a new genomic organization of chickpea *pgip* genes (Figure 1). Full-length cDNA size (bp), ORF size (bp), predicted protein size (aa), predicted signal peptide size (aa), theoretical molecular mass (kDa), and pI for all four *Capgips* are presented in Table 1.

**FIGURE 1:**
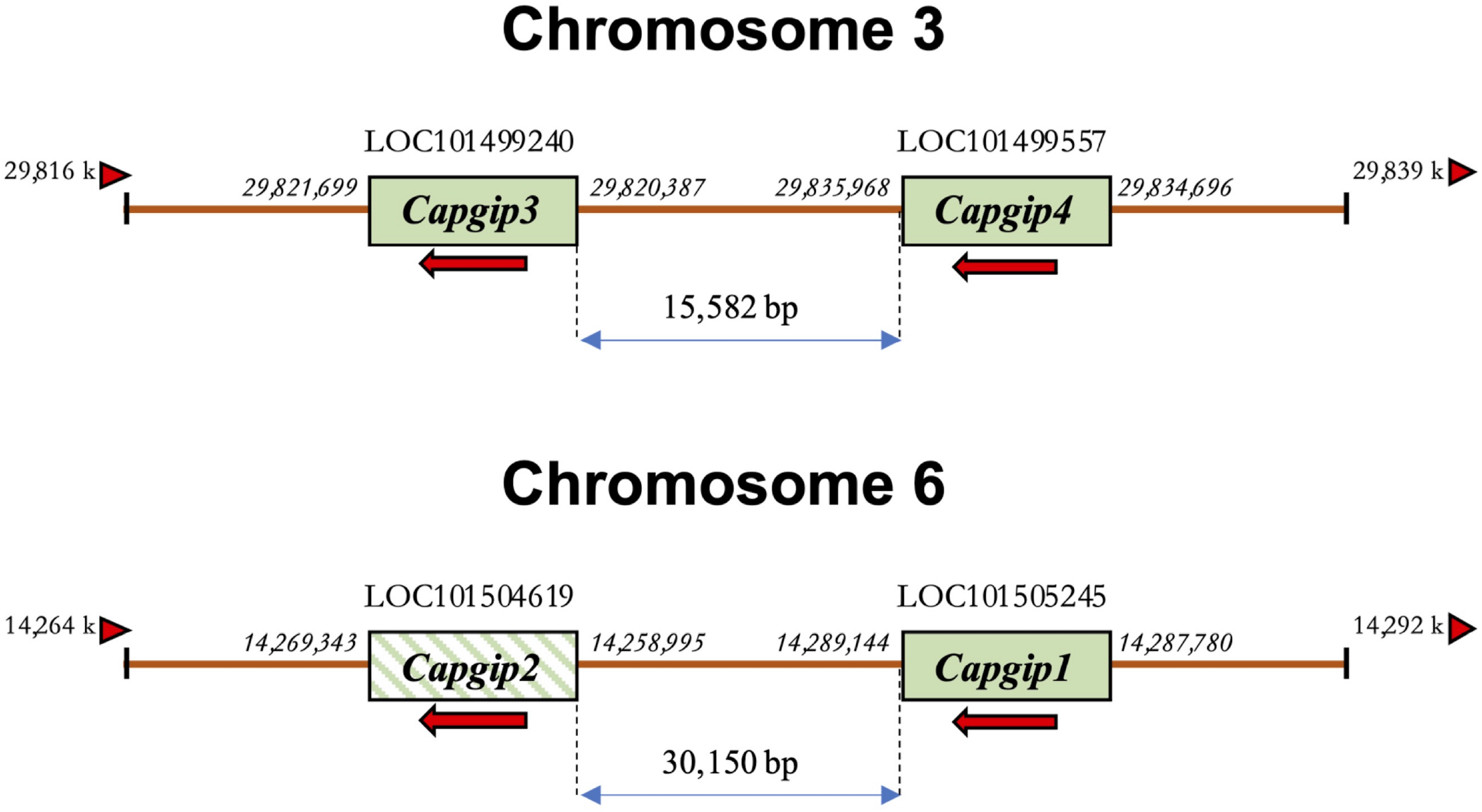
Schematic representation of the revised genomic organization of the *pgip* gene family in *Cicer arietinum* showing two loci for *Capgips*, chromosomes 3 and 6. The numbers on the side of genes represent their start and stop codon location on the genome. The numbers between the genes denote distances in base pairs. Arrows indicate the transcriptional direction of coding regions. The arrowhead triangles indicate the orientation/direction of the DNA strand on which the genes are placed. Crossed box for *Capgip2* indicates the presence of an intron within the gene.

Sequence analysis was conducted for all four CaPGIPs. Predicted proteins of CaPGIP1, CaPGIP3, and CaPGIP4 exhibited a typical PGIP sequence identity. SignalP 5.0 (Armenteros et al., 2017) projected 36, 22, and 20 amino acid signal peptides for CaPGIP1, CaPGIP3, and CaPGIP4, respectively (Table 1). As illustrated in Figure 2, the predicted mature protein sequences for CaPGIP1, CaPGIP3, and CaPGIP4 featured an N-terminal domain, a central LRR domain, and a C-terminal domain. Like many PGIPs, the central domain of CaPGIP1, CaPGIP3, and CaPGIP4 is comprised of 10 imperfect leucine-rich repeats (LRRs). Each is about 24 amino acids long and perfectly matches the extracytoplasmic LRR (eLRRs) consensus sequence xxLxLxx.NxLx..GxIPxxLxxL.xxL (Di Matteo et al., 2003). These tandemly repeated LRRs fold into a characteristic curved and elongated PGIP shape (Maulik et al., 2009), as observed in homology 3D models generated using PvPGIP2 as a template for CaPGIP1, CaPGIP3, and CaPGIP4 (Figure 3). The secondary and tertiary structures of CaPGIP1, CaPGIP3, and CaPGIP4 indicate that all 10 LRRs contain an β-turn motif (xxLxLxx) that folds into β1 sheets. β1 sheets occupy the PGIP scaffold’s inner concave face, which is the site for PG interaction. Aside from that, all LRRs have β2 sheets and 310-helixes. PGIPs are glycoproteins with N-glycosylation sites ((N–x–S/T; where N is asparagine, x can be any amino acid except proline (P), S is serine, and T is threonine). As a result, the NetOGlyc 4.0 server predicted five, three, and eight N-glycosylation sites in CaPGIP1, CaPGIP3, and CaPGIP4 proteins, respectively. CaPGIP1, CaPGIP3, and CaPGIP4 proteins all contain conserved cysteine residues that form disulfide bridges that are essential for the maintenance of secondary PGIP structures (Veronico et al., 2011). CaPGIP1 has eight, CaPGIP3 has ten, and CaPGIP4 has nine cysteine residues.

**FIGURE 2:**
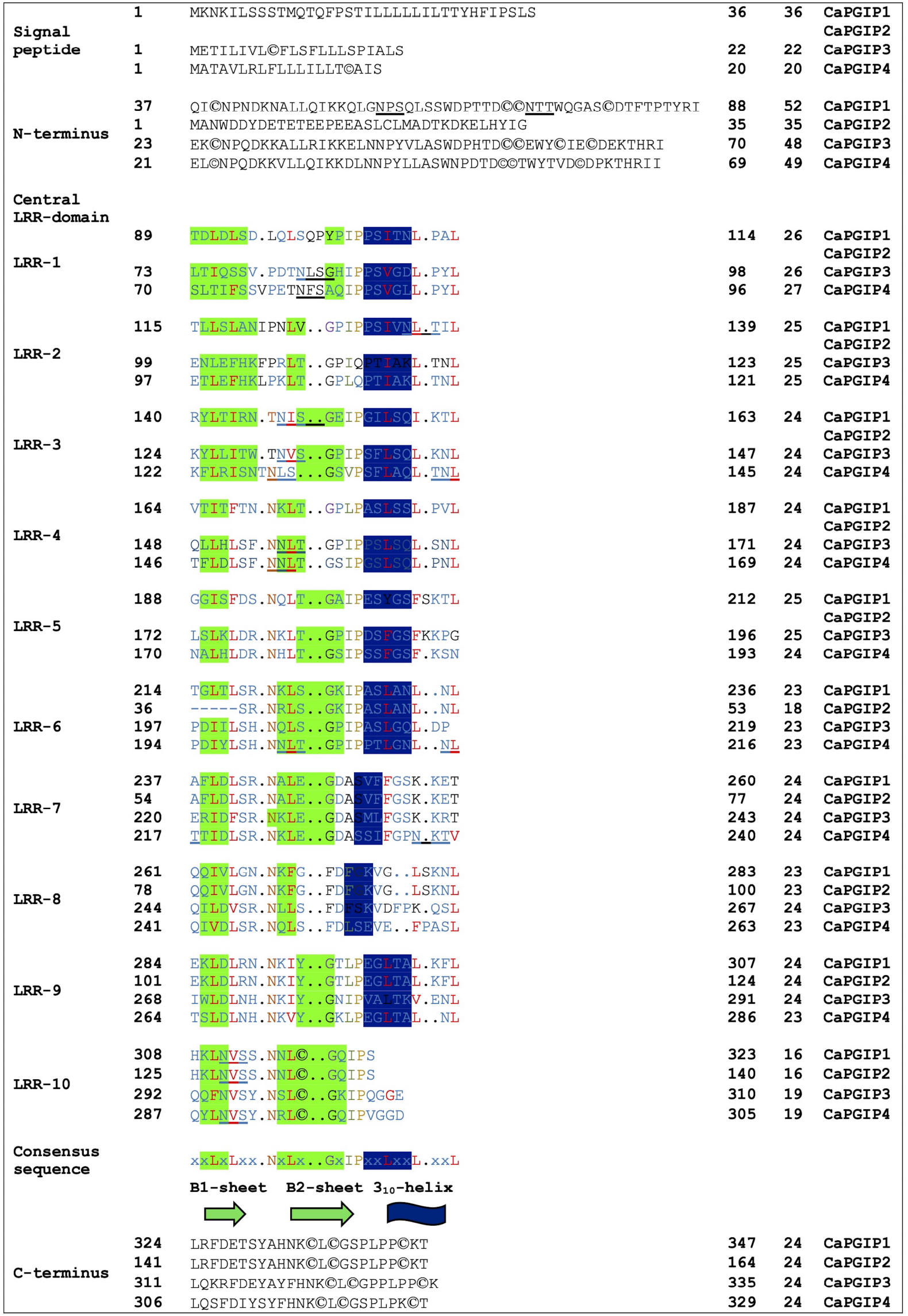
Translated structure of CaPGIPÍ, CaPGIP2, CaPGIP3, and CaPGIP4 based on PvPGIP2. A) Signal peptide, B) N-terminal domain, C) central LRR domain and D) C-terminal domains are indicated. Secondary structure elements (sheets β 1, β 2, and 3io-helix) are highlighted. Five N-glycosylation sites (N-X-S/T) are underlined, cysteine residues are encircled.

Interestingly, CaPGIP2, on the contrary, lacked many of the above sequence identities. The absence of signal peptide suggested it is not a secretory protein. Its secondary structure reveals that it lacked more than half of the LRR modules, with just the 6^th^ to 10^th^ LRRs. This short LRR sequence on CaPGIP2’s C-terminal perfectly matches CaPGIP1’s C-terminal. The CaPGIP2 homology 3D model indicated the absence of the distinctive concave face that harbors PG interaction sites (Figure 3). CaPGIP2 has one N-glycosylation site and five cysteine residues, fewer than its counterparts. All these observations imply that CaPGIP2 might be mischaracterized as a PGIP gene and can be a pseudogene and not a PGIP, lacking the typical characteristics a PGIP and therefore likely nonfunctional.

**FIGURE 3:**
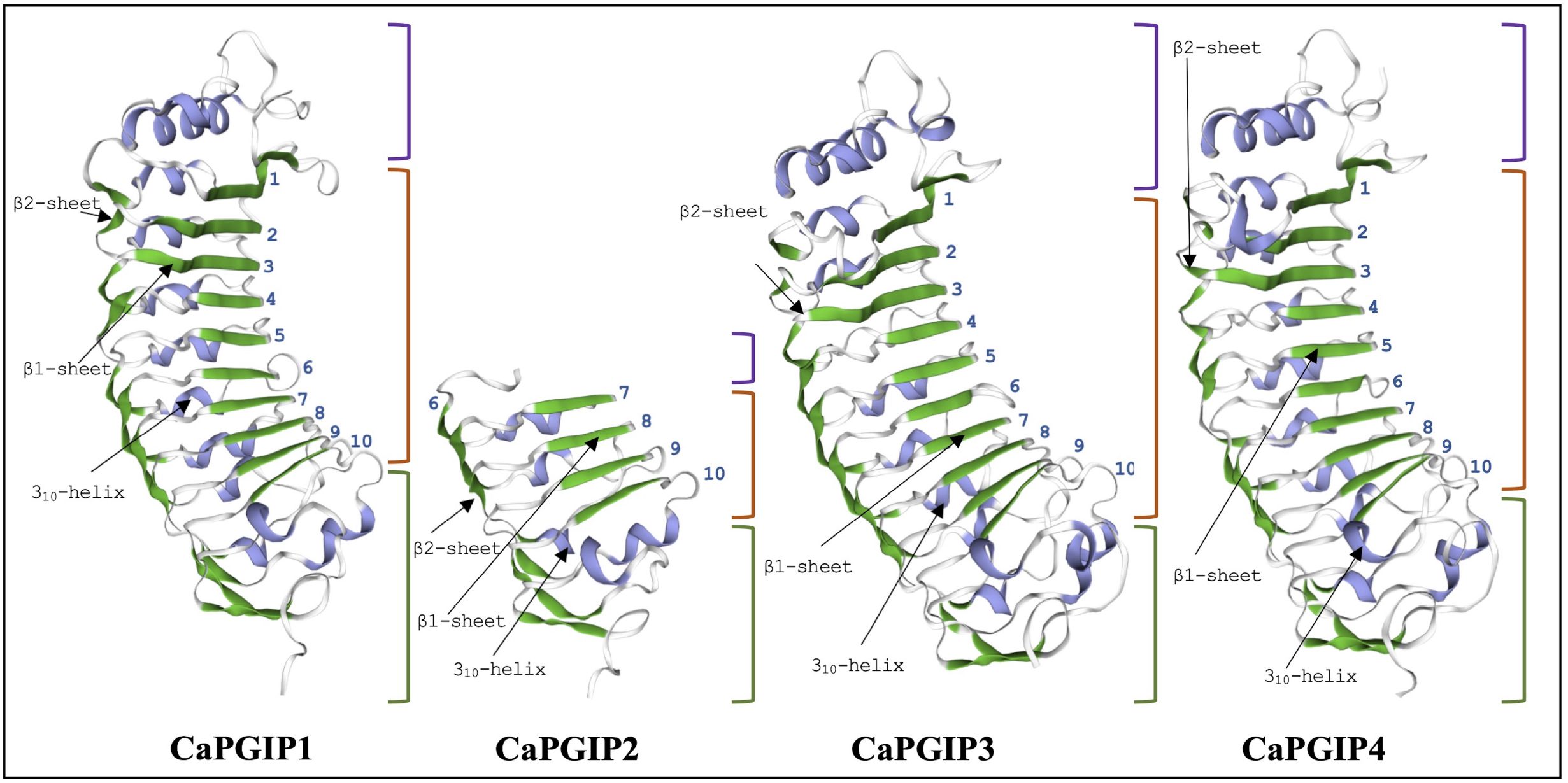
Homology 3D model of CaPGPl, CaPGIP2, CaPGIP3, and CaPGIP4 using PvPGIP2 as a template, β 1, and β 2 sheets are indicated by green color. 3_10_-helixes are indicated with purple color. Purple-colored brackets indicate N-terminal domains, orange-colored brackets indicate central LRR domain with LRR numbers marked and green-colored brackets indicate C-terminal domain.

### 3.2 Sequence comparison and phylogenetic analysis of the CaPGIP proteins

Multiple sequence alignment showed that CaPGIP amino acid sequences are highly similar to those of other legumes such as soybean, common bean, runner bean, tepary bean, lima bean, barrel clover, peas, mung bean, and alfalfa, with the presence of five conserved cysteine residues shared by all. The higher similarity was observed in the β2-sheet regions, along with variable portions present in both β-sheets, as evident in plant-specific LRR proteins (Figure 4). This variability is most likely responsible for the presence of multiple recognition specificities to target broader pathogen PGs (De Lorenzo et al., 2001, and Matsushima and Miyashita, 2012). This sequence alignment demonstrates that PGIPs are highly conserved within the legume family.

**FIGURE 4:**
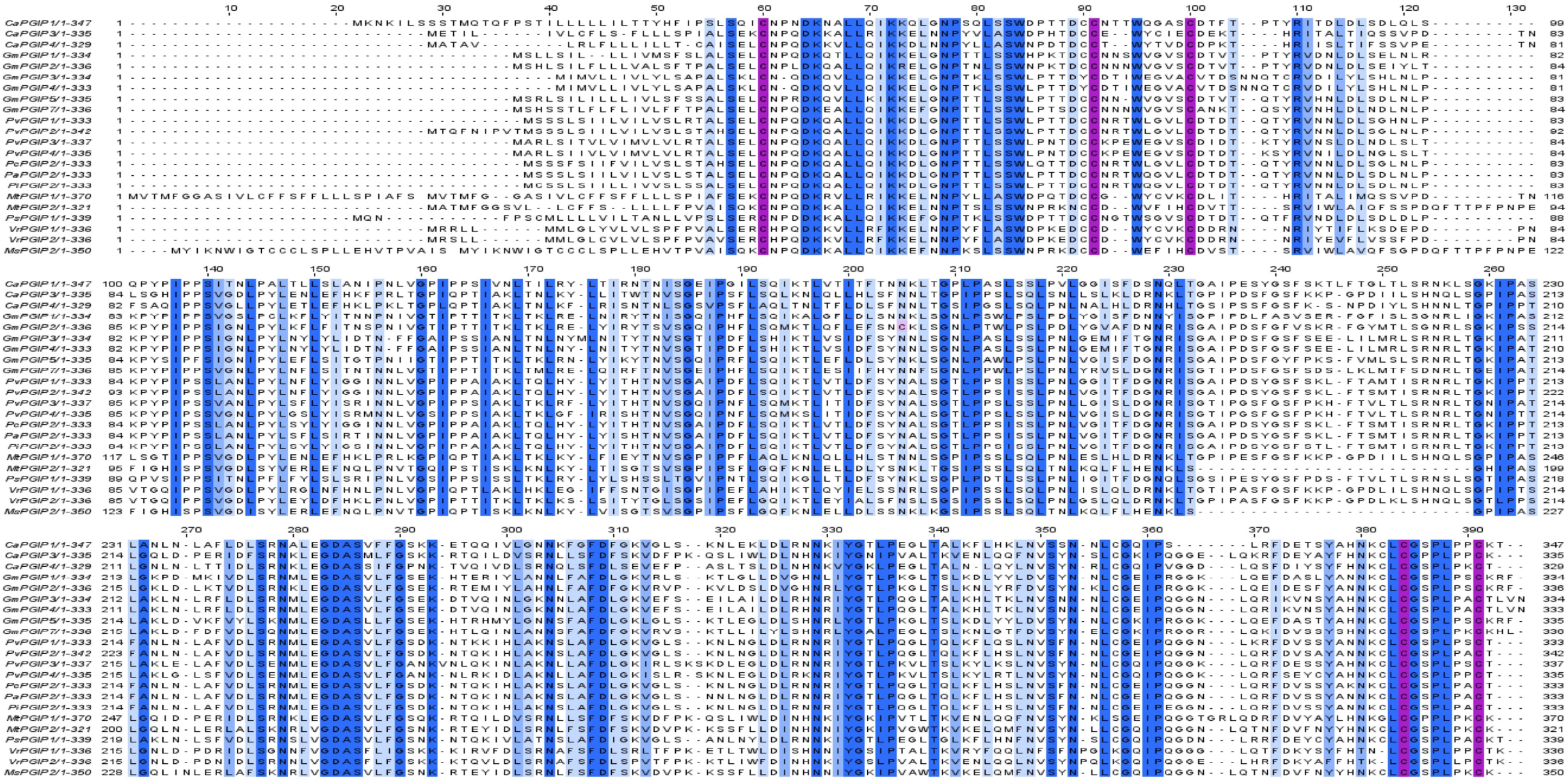
Multiple sequence alignment of chickpeas PGIPs *[Capgip1* (XP_004504732.1), *Capgip3* (XP_004493557.1), *Capgip4* (XP_01256925 8.1)], with other legume PGIPs such as soybean [GmPGIPl(XP_003524772.1), GmPGIP2 (CAI99393.1), GmPGIP3 (NP_OO 1304551.2), GmPGIP4 (CAI99395.1), GmPGIP5 (XP_003524769.1), GmPGIP7 (XP_OO3531070.1)], common bean [PvPGIPl(CAIl 1357.1), PvPGIP2 (P58822.10, PvPGIP3 (CAI11359.1), PvPGIP4 (CAI11360.1)], runner bean [PcPGIP2 (CAR92534.1)], tepary bean [PaPGIP2 (CAR92533.1)], lima bean [PiPGIP2 (CAR92537.1)], barrel clover [MtPGIP1 (XP_003625218.1), MtPGIP2 (XP_024626259.1)], peas [PsPGIPl (AJ749705.1)], mung bean [VrPGIPl (ATN23902.1), VrPGIP2 (ATN23895.1), and alfalfa [MsPGIP2 (ALX18673.1)]. Blue color bar indicates conservation more than 50%, brighter the bar more the conservation, pink color bar indicates cysteine residues.

In the phylogenetic analysis (Figure 5), CaPGIPs were compared to 45 other known PGIPs from various crop families. The tree (Figure 5) comprises five main branches that are separated into monocots and dicots. The Poaceae family is represented by one cluster, the majority of the legume PGIPs are represented by a second cluster, three PGIPs from *Beta vulgaris* are represented by a third cluster. CaPGIP3, and PGIPs from *Vigna radiata* are represented by a fourth cluster, and the remaining PGIPs from other families, such as Actinidiaceae, Apocynaceae, Brassicaceae, Caricaceae, Cucurbitaceae, and Malvaceae, form the final cluster. CaPGIP1 and CaPGIP2 are members of the Leguminosae cluster and exhibit significant similarities to pea PGIP, PsPGIP1. CaPGIP3 and CaPGIP4 are outside the Leguminosae cluster, with CaPGIP3 sharing a high degree of similarity with its other legume counterparts, *Vigna radiata* PGIPs, VrPGIP1, and VrPGIP2. CaPGIP4 is in a separate cluster that includes PGIPs from various plant families. This suggests that CaPGIP1 and CaPGIP2 may have characteristics and functions like legume PGIPs, whereas PGIP3 and CaPGIP4 may have characteristics and functions similar to PGIPs belonging to non-legume families.

**FIGURE 5:**
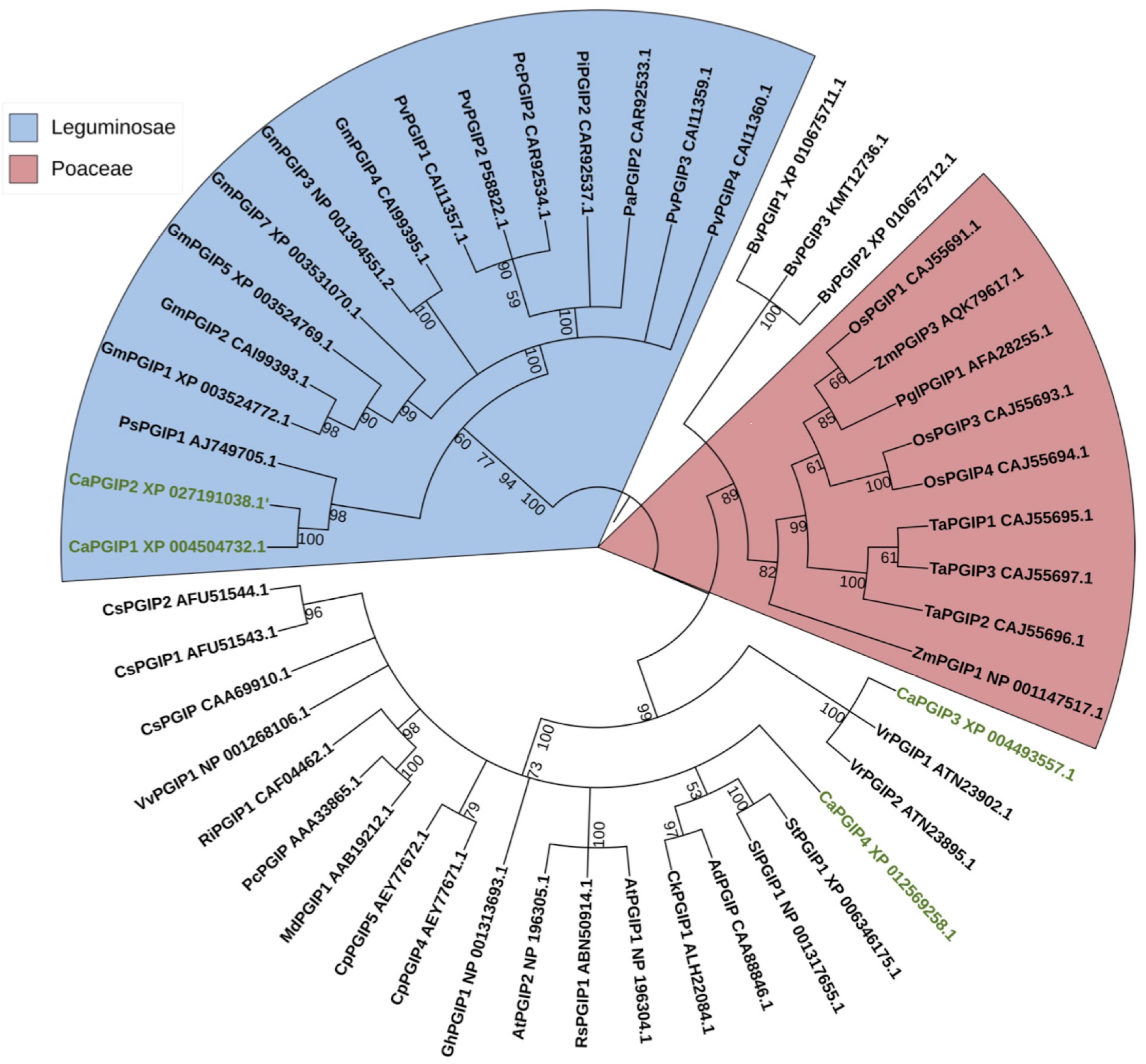
Phylogenetic tree illustrating the relationship between several PGIPs from various crop families and CaPGIPs. CaPGIPs are highlighted in green. The values on the branches correspond to bootstrap values. *Cucumis sativus* PGIPs (CsPGIPs) are used as outgroup.

### 3.3 Promoter analysis of *Capgips*

To locate regulatory DNA elements, the 1,500 bp upstream sequence for all *Capgips* was analyzed. The TATA box and CAAT box motifs, which are widely found in functional gene promoter and enhancer regions (Svensson et al., 2006, and Xue et al., 2002), were discovered close to the start codons. CAAT and TATA box sequences were found at −40 and −229 upstream of *Capgip1*, respectively, and at −23 and −55 upstream of *Capgip3*. The TATA box was located at a −30 position in the *Capgip4* upstream sequence, whereas CAAT was at a −37 position. Elements/motifs associated with plant responses to hormones such as abscisic acid, gibberellic acid, jasmonic acid, and salicylic acid were also identified. Motifs for wounding response were identified as well. Crucially, numerous elements associated with pathogenicity responses were identified in the promoter regions of the *Capgips*. Table 2 lists these putative cis-acting regulatory elements, as well as their locations and roles. Aside from the motifs mentioned above, additional cis-elements known to mediate tissue-specific activity and plant physiological processes were identified. Stress-related cis-acting regulatory elements associated with drought, dehydration, water, high light, and low-temperature stress, were also identified. All these elements are listed in Supplementary Table 2.

### 3.4 Cloning and characterization of *Capgips*

*Capgip* genes were cloned and sequenced from the chickpea cultivar “Dwelley”. *Capgip1* sequence matched GenBank sequence XM 004504675. However, a C was replaced by a T at the 720^th^ nucleotide position, which was a synonymous substitution with no change in the coded amino acid. *Capgip3* sequence matched the GenBank sequence XM 004493500, and *Capgip4* sequence matched the GenBank sequence XM 012713804. Transcripts for *Capgip2* could not be amplified even with different sets of primers; hence, all subsequent investigations focused on *Capgip1*, *Capgip3*, and *Capgip4*.

### 3.5 Subcellular localization of *Capgips*

DeepLoc-1.0 uses sequencing information to predict the subcellular localization of plant proteins. Based on the presence of signal peptides, it was inferred that *Capgip1, Capgip3*, and *Capgip4* were secretory and classified as extracellular proteins. *Capgip2*, on the other hand, was predicted to be found in the mitochondrion and cytoplasm. To validate those predictions, *Agrobacterium* cells carrying binary vectors of *Capgips* and GFP fusions were infiltrated into *N. benthamiana* leaves for transient expression of encoded proteins in leaf mesophyll and epidermal cells. According to the excitation curves, the fluorescence of *Capgips-GFP* fusion proteins was the same as GFP. *Capgip1, Capgip3*, and *Capgip4* fluorescence were visible on the cell boundaries (Figure 6). As a result, they were most likely found on the cell wall or plasma membrane. *Capgip2-GFP* fluorescence was seen inside the cell in the cytoplasm and ER, and *Capgip2* was most likely found in the cytoplasm and ER (Figure 6).

**FIGURE 6:**
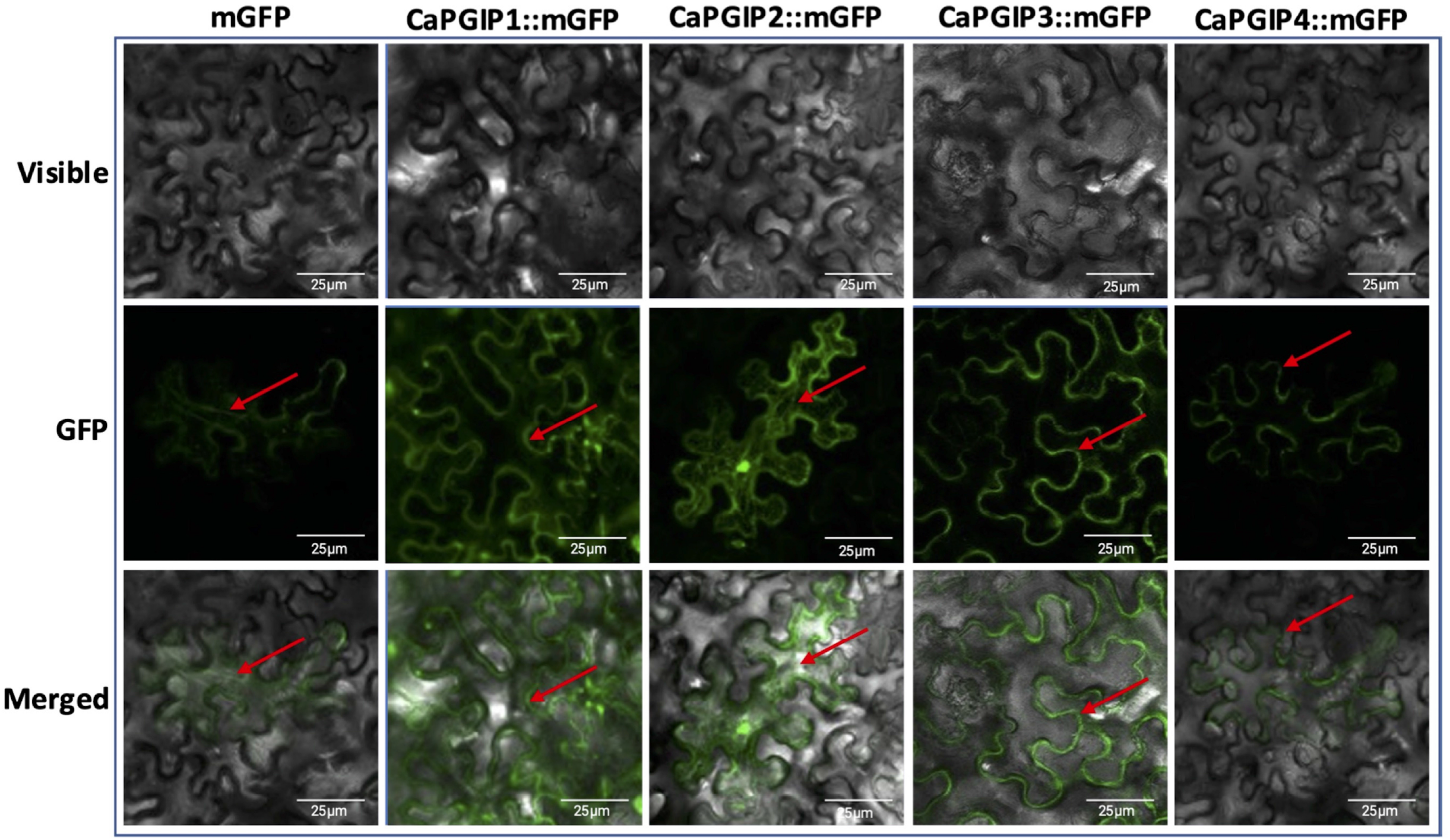
Subcellular localization of *Capgips*. Full-length *Capgips* fused with a green fluorescent protein (GFP) are transiently expressed in *Nicotiana benthamiana* leaves by agroinfiltration. The images show the fluorescence emitted by fusion proteins was captured 72 hours after infiltration using a laser scanning confocal microscope as mGFP fluorescence in green color, visible light in brightfield images, merged as merged images of mGFP and visible. Cells transformed with mGFP is the control. Red arrows indicate localization

### 3.6 Absolute quantification of *Capgips* transcripts to understand basal gene expression levels

*Capgip* transcript levels were investigated at four growth stages using the indeterminate type of Kabuli chickpea variety Dwelley, which matures in 110 to 120 days. Based on the timing of infection by the major chickpea fungal pathogens, four growth stages were selected (Bretag and Horsham, 2004; Jiménez-Fernández et al., 2011; Khan et al., 2012; Knights and Hobson, 2016; Markel et al., 2008; Mazur et al., 2002; Moore et al., 2011; West et al., 2003; Wunsch, 2014). RT-qPCR was utilized to determine the absolute *Capgips* expression levels at various chickpea “Dwelley” growth stages (Table 3). Transcripts for *Capgip1, Capgip3*, and *Capgip4* were ubiquitously detected in all the studied tissues. In the vegetative stages V1 and V6 stages were investigated (Figure 8). *Capgip1, Capgip3*, and *Capgip4* transcript levels were higher in the leaf at the V1 stage. For *Capgip1* and *Capgip3*, they were expressed the most in the leaves, followed by the root and then the stem. The highest expression of *Capgip4* was found in the leaves, stems, and roots. In the V6 vegetative growth stage, higher amounts of gene transcripts for *Capgip1, Capgip3*, and *Capgip4* were found in stem tissues, in contrast to the V1 stage, where stem transcript levels were lower. For *Capgip3* and *Capgip4*, the stems are expressed the most, followed by the root and then the leaves. For *Capgip1* after stems highest expression was seen in leaf and least in the root.

**FIGURE 7:**
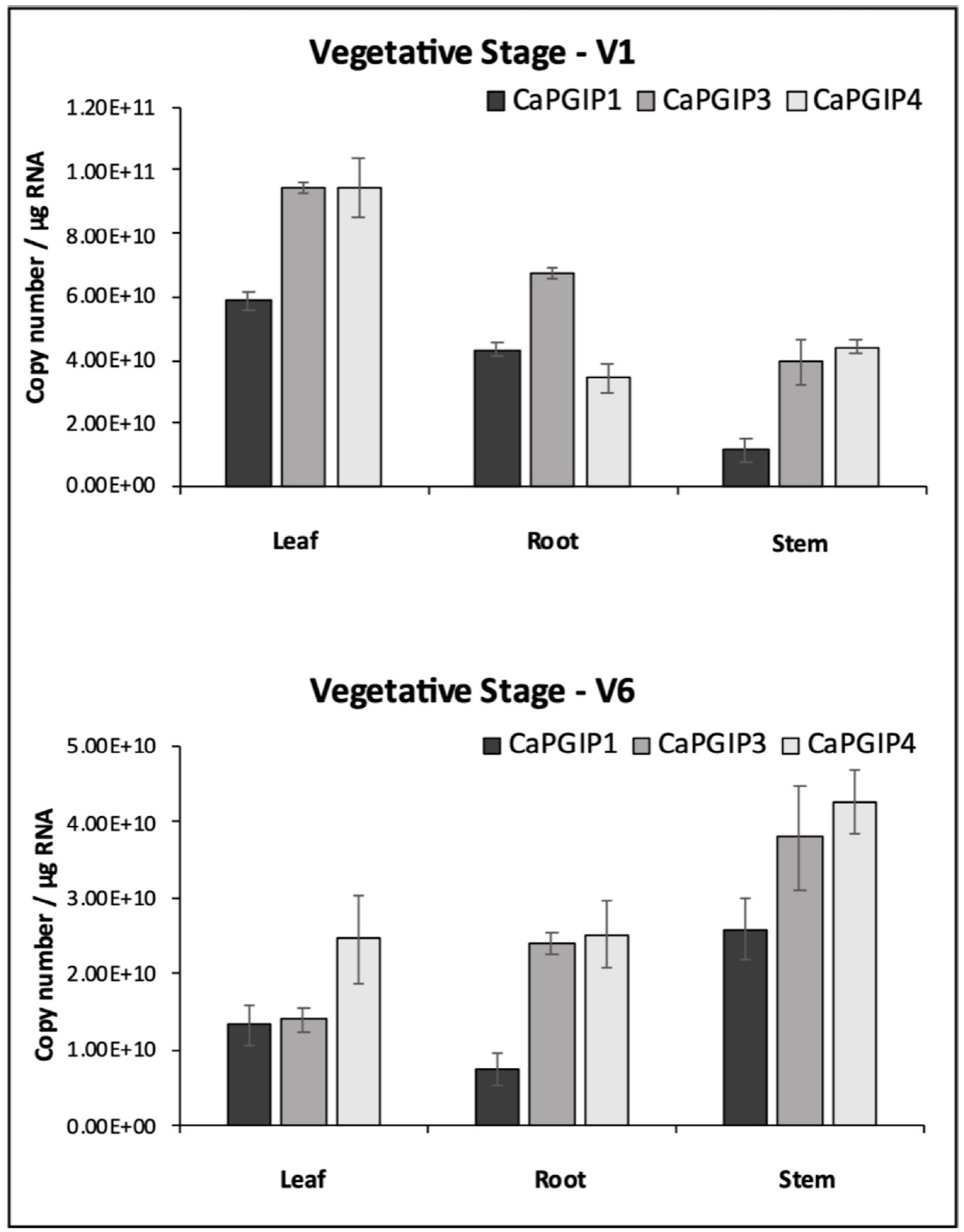
Absolute expression of *Capgips*. The number of transcript copies of *Capgip1, Capgip3*, and *Capgip4* was calculated in the leaf, stem, and root at vegetative growth stages VI and V6. The abundance was normalized by the amount of internal control 18 SrRNA and 25 rRNA. The values are the means of three biological replicates each with three technical replicates. Error bars indicate standard deviation among the biological replicates (N =3).

**FIGURE 8:**
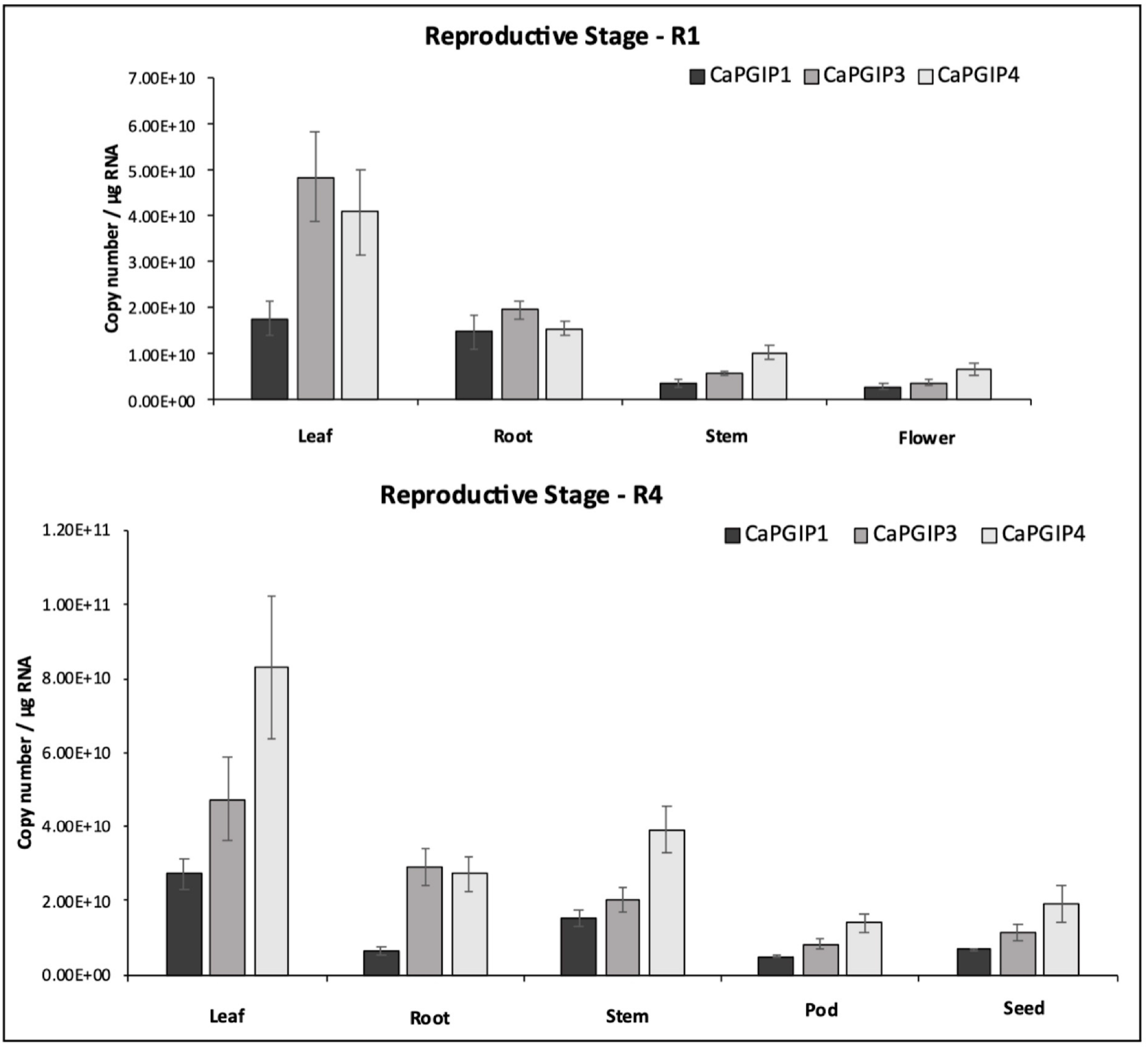
Absolute expression of *Capgips*. The numbers of transcript copies of *Capgip1, Capgip3*, and *Capgip4* were calculated in the leaf, stem, root and flower at reproductive growth stage R1 and leaf, stem, root, pod and seed at reproductive growth stage R4. The abundance was normalized by the amount of internal control 18 SrRNA and 25 rRNA. The values are the means of three biological replicates each with three technical replicates. Error bars indicate standard deviation among the biological replicates (N = 3).

In the reproductive stages, the transcript levels were analyzed for flower, pod, and seed along with leaf, stem, and root (Figure 9). Similar to R1, leaves showed the highest expression for all the *Capgips* in R1. The roots had the highest expression for *Capgips* after the leaves, followed by the stems, and the least in the flowers. In the reproductive stage, R4 gene transcripts for all *Capgips* were found highest in leaves and the least in pods. Apart from leaves and pods, for *Capgip1*, higher expression was in stems, followed by seeds, then roots. Gene transcripts for *Capgip3* were higher in roots, then stems, and then seeds. Gene transcripts for *Capgip4* were higher in stems, then roots, and then seeds. Overall, leaves displayed the highest expression for all *Capgips*, and *Capgip 4* showed the highest expression, which was about one and a half times the expression of *Capgip3* and double the expression of *Capgip1*.

## 4 Discussion

Several plant species have PGIP-encoding small gene families; these multigene families encode proteins with similar LRR domains but different PG-inhibitory capabilities (D’Ovidio et al., 2006). This study demonstrated that the chickpea genome has a *pgip* family of potentially three functional genes (*Capgip1, Capgip3*, and *Capgip4*) that are present on two chromosomes. Soybean and barrel clover are the only two other legumes where *pgips* are located on multiple chromosome (D’Ovidio et al., 2006, and Song and Nam, 2005). Like the genomic distribution of *Capgips*, legume *pgips* are distributed across a broader genomic region, as observed in soybean, where *Gmpgip1, Gmpgip2*, and *Gmpgip5* are located on chromosome 5, and span a ~ 19.5 kbp region, while *Gmpgip3, Gmpgip4*, and *Gmpgip7*, present on chromosome 8, span a ~ 21 kbp region. In common bean, *Pvpgip1, Pvpgip2, Pvpgip3*, and *Pvpgip4* span an area of ~ 50 kbp on chromosome 2 (D’Ovidio et al. 2004, and D’Ovidio et al. 2006).

So far, over 18 legume PGIPs have been characterized in 9 species. The reported peptide size of these legume PGIPs ranged from 321 amino acids for MtPGIP2 to 342 amino acids for MtPGIP1. All these PGIPs contain signal peptides, with GmPGIP4 having the shortest with 17 amino acids and PvPGIP2 having the longest with 29 amino acids. Protein molecular mass (kDa) ranged from 35.92 for MtPGIP2 to 38.17 for MtPGIP1. And the isoelectric point (pI) ranged from 6.79 for PsPGIP1 to 9.48 for PvPGIP4. Except for MsPGIP2 of alfalfa, all other legume PGIPs had 10 LRRs and all were intronless. Only MsPGIP2’s genomic sequence contains 9 LRRs and a 154-bp intron. CaPGIP1, CaPGIP3, and CaPGIP4 exhibit very similar above-mentioned characteristics, with the exception that CaPGIP1 has the longest signal peptide among the known legume PGIPs so far, with 36 amino acids. Because of selection for transcription efficiency, conserved genes with relatively high levels of expression tend to lose introns (Zou et al., 2011). Intronless genes such as PGIP may play key roles in plant growth, development, or response to biotic or abiotic stresses (Liu et al., 2021).

Like many legume PGIPs, CaPGIPs possessed LRRs matching the extracytoplasmic sequence, eLRRs are important in plant defense as they function as receptor-like proteins or receptor-like kinases to recognize diverse pathogen molecules, and plant hormones (van der Hoorn et al., 2005). The presence of xxLxLxxNxL core consensus in eLRRs is responsible β – sheet structure formation (Zambounis et al., 2012). The LRRs are arranged tandemly in the PGIP’s distinctive curved and elongated shape. The β-sheets are parallelly organized on the inner side of the protein to form the concave face, while the 310-helixes are parallelly organized on the outer side to make the convex face (Kobe and Deisenhofer, 1993). Solvent-exposed residues on the concave-sheet surface bind to pathogen molecules. CaPGIPs, have two types of β– sheets (β1 and β2) and 3_10_-helixes. All LRRs of plant defense proteins have β1 sheet; however, the presence of β2 sheets is unique and seen only in PGIPs (Di Matteo et al., 2003). CaPGIPs are predicted to have several N-glycosylation on these β-sheets which are vital for ligand binding for disease resistance, and the heterogeneity in β-sheet residues or the glycosylation patterns contributes to the varying recognition specificities of LRR proteins (Ramanathan et al., 1997, and van der Hoorn et al., 2005). The deduced CaPGIP proteins also contain conserved cysteine residues that form disulfide bridges crucial for the maintenance of secondary structures in PGIP (Veronico et al., 2011).

Previous research indicated that truncated variants of functioning NBS-LRR genes can be found within 100 kb of fully functional NBS-LRR genes. These truncated genes are often pseudogenes because of alternative splicing. These pseudogenes have large deletions due to various transposition events (Marone et al., 2013). CaPGIP2 is found within 30 kb of CaPGIP1. CaPGIP2’s C-terminal end matches CaPGIP1 perfectly, implying that the majority of the central LRR domain and N-terminus might be deleted. Also, the presence of a nearly 10-kbp intron is atypical for PGIPs and most functional genes. Since it also lacks a fundamental signal peptide, it is classified as a non-secretory protein. For these reasons, CaPGIP2 is not considered a true PGIP in this study.

Multiple sequence alignment of CaPGIP deduced amino acid sequences with various other legume PGIPs revealed that CaPGIP amino acid sequences are highly similar to those of other characterized, suggesting that CaPGIPs, like PGIPs from other legumes, are highly conserved. Interestingly, alignment also indicated that legume PGIPs are highly conserved at the β2-sheet sites, which may be since they are unique only to PGIPs. As per the phylogenetic analysis, CaPGIPs have a high degree of similarity with PGIPs from different plant sources. CaPGIP1 shared the most similarities of 70% with *Pisum sativum* PsPGIP1 (AJI49705.1), and 69 % with *Glycine max* GmPGIP3 (NP 001304551.2). PsPGIP1 has been identified as a possible protective factor against the pea-cyst nematode *Heterodera goettingiana* (Veronico et al., 2011). Encoded protein products of GmPGIP3 inhibited Sclerotinia sclerotiorum PGa, *Sclerotinia sclerotiorum* PGb, and crude PG extracts of *Fusarium moniliforme, Botrytis aclada, Aspergillus niger, Botrytis cinerea, Colletotrichum acutatum, and Fusarium graminearum*. Transgenic wheat expressing GmPGIP3 showed enhanced resistance to *Gaeumannomyces graminis* var. *tritici*, *Bipolaris sorokiniana* (D’Ovidio et al., 2006, and Wang et al., 2015). These reports suggest that CaPGIP1 may have similar functions as PsPGIP1 and GmPGIP3 and might be involved in disease inhibition of nematodes or fungi. CaPGIP1 may have functions similar to PsPGIP1 and GmPGIP3 and may be involved in nematode or fungal disease inhibition. CaPGIP3 is more similar to the two tightly linked PGIPs of *Vigna radiata*, with 64% and 69% similarity to VrPGIP1 and VrPGIP2, respectively. Both VrPGIPs are known to provide resistance to bruchids (*Callosobruchus spp*) (Kaewwongwal et al., 2017), and CaPGIP3 may play a similar role in chickpeas against bruchids. CaPGIP4 had higher similarity to tree fruit PGIPs, with 72 % similarity to *Malus domestica* MdPGIP1 (AAB19212.1) and 71 % similarity to *Pyrus communis* PcPGIP (AAA33865.1). MdPGIP1 protein inhibited PG production in *Colletotrichum lupini* and *Aspergillus niger* (Oelofse et al., 2006). Unlike CaPGIP1 and CaPGIP3, CaPGIP4 may only be effective against pathogenic fungi.

TATA boxes function as a motif for recruiting transcription initiation machinery and RNA polymerase II, while CAAT boxes improve protein binding (Joubert et al., 2013, and Liao et al., 2015). TATA and CAAT boxes are conserved eukaryotic cis-elements that are found in many plant gene promoters, including *pgips. Capgip1, Capgip3*, and *Capgip4* all have several TATA and CAAT boxes upstream of the start codon ATG, indicating that they are functioning genes. Plant *pgips* are typically expressed after pathogen infection and wounding response (Kalunke et al., 2015), hence the presence of multiple pathogenicity-related and wounding motifs in the *Capgips* promoter region. Apart from pathogens, *pgips* are triggered by phytohormone treatment in several plant species (Stotz et al. 1993; Yao et al. 1999, and Ferrari et al. 2003). *Pgip* expression in rice, alfalfa, and pepper is induced by abscisic treatment. *pgips* from rapeseed, rice, barrel clover, and pepper are triggered by jasmonic acid and salicylic acid (Hegedus et al., 2008, Janni et al, 2006, Lu et al, 2012, Song and Nam 2005, and Wang et al., 2013). Rice *pgips* are induced by gibberellic acid treatment (Janni et al, 2006, and Lu et al, 2012). The presence of multiple cis-acting elements in *Capgips* that regulate abscisic acid, gibberellic acid, jasmonic acid, and salicylic acid pathways suggests that they may play comparable roles to those seen in many plant *pgips*. Like other families of defense-related genes*, pgips* demonstrate tissue-specific activity. The grapevine *Vvpgip1* gene is only expressed in roots and ripening berries, and its expression is developmentally regulated (Joubert et al., 2006). Several cis-acting elements that influence tissue-specific responses, particularly root-specific responses, were found in all *Capgips*, indicating a role in plant development or resistance to pathogens that enter the plant system through the roots. *B. juncea pgips* are associated with high temperature and drought stresses (Bhardwaj et al., 2015), while Arabidopsis *Atpgip1*, and apple’s *Mdpgip1*, are induced in response to cold stress (Kalunke et al., 2015). The presence of regulatory elements associated with drought, dehydration, water, high light, and low-temperature stress in the *Capgip* promoter suggests that they may play a role in plant stress.

Bioinformatic analysis and subcellular localization confirmed that CaPGIP1, CaPGIP3, and CaPGIP4 are secretory proteins, and they are located in the plasma membrane or the cell wall. DeepLoc-1.0 analysis indicated CaPGIP2 might be found in the mitochondrion or the cytoplasm, and localization experiments revealed *Capgip2* was found in the cytoplasm and ER. The locations of CaPGIPs within the plasma membrane/cell wall were consistent with the locations of other legume PGIPs. Localization on the plasma membrane is crucial as these proteins play a role in defense responses as cell surface receptors to detect pathogen PGs in the apoplast (Rodriguez-Palenzuela et al., 1991). CaPGIP2 localization to the cytoplasm and ER might be because of the lack of signal peptide in CaPGIP2 and suggests it might not be involved with the PG interaction in the apoplast.

Several studies investigated *pgip* gene expression in response to external stimuli. Analyzing constitutive expression in similar environmental conditions, on the other hand, allows for the correlation of the expression of various pgips within a crop. Furthermore, pathogens can infect plants at any stage of their life cycle, and *pgip* families, like other defense-related gene families, have been demonstrated to exhibit variable constitutive expression patterns (Kalunke et al., 2015). Because *pgips* exhibit functional redundancy and sub-functionalization at the protein level (De Lorenzo et al., 2001; Federici et al., 2006), tissue-specific expression of PGIPs is feasible, allowing pgips to respond more effectively to a variety of environmental stimuli (D’Ovidio et al., 2004). Increased transcript abundance of *pgips* during fungal infection enhances the likelihood of plant defense. In terms of pathogen PG specificity, plant PGIPs can express at higher levels in distinct growth stages and tissues that correspond to pathogen infection (Cantu et al., 2008). Absolute expression analysis showed *Capgips* has higher expression in leaf tissue and the least in pod tissues. *B. vulgaris’s Bvpgip* genes were reported to be highly expressed in roots in comparison to leaf tissue during normal growth and development (Li and Smigocki, 2016). *Carica papaya’s Cppgip4* and *Cppgip6* were shown to be ubiquitously expressed in root, stem, leaf, seed fruit pulp and peel. However, the *Cppgip* transcripts were most abundant in fruit pulp and peel and decreased during ripening (Broetto et al., 2015). Tissue-specific differences have been reported in apples, where higher transcript abundance was in leaves and fruit, least in the stem (Zhang et al., 2010). In blackberry *pgip* gene expression was more abundant in young leaves and fruit compared to old leaves and ripe fruit (Hu et al., 2012). In raspberries, *pgip* transcripts were detected in fruit but not in flowers (Johnston et al., 1993). The varying expression levels of *Capgips* in different tissues at different growth stages indicates that *Capgips* might respond to different stimuli.

## 5 Conclusion

In conclusion, this study is the first to characterize chickpea PGIPs. Two additional PGIPs on chromosome 3, CaPGIP3 and CaPGIP4 were identified in addition to the previously reported CaPGIP1 and CaPGIP2 on chromosome 6. CaPGIP1, CaPGIP3, and CaPGIP4 displayed a typical PGIP sequence identity with an N-terminal domain, a central LRR domain with ten imperfect LRRs, and a C-terminal domain. Multiple sequence alignment shows that CaPGIP amino acid sequences are highly like those of other described legumes. The phylogenetic study of CaPGIPs indicated that CaPGIP1 and CaPGIP3 are closer to legume PGIPs, and CaPGIP4 falls outside the legume PGIP cluster. *Capgip’s* promoter sequences harbor cis-elements that regulate response to various external stimuli. CaPGIP1, CaPGIP3, and CaPGIP4 are localized to the cell wall or plasm membrane. Absolute quantification of the *Capgip* transcript levels under normal conditions demonstrates that *Capgips* have constitutive tissue-specific expression. Interestingly, every experiment indicated CaPGIP2 could not be a true PGIP, and this necessitates modifying the genomic organization of CaPGIP. All these findings indicate that CaPGIPs could have similar functions as other crop PGIPs, and additional research is needed to determine CaPGIPs’ roles against pathogen PGs.

## 6 Author Contributions

VE and WC conceived the study and designed the experiments. VE performed the experiments. VE analyzed the data. VE wrote the manuscript with contributions from all co-authors. All authors read and approved the final manuscript.

## 7 Acknowledgments

We would like to thank Dan Mullendore of Franceschi Microscopy and Imaging Center, Washington State University, Pullman, WA for assistance with confocal microscopy. We would also like to thank Sheri McGrew, Grain Legume Genetics Physiology Research, USDA ARS, Pullman, WA for assistance with maintaining plants in the green house. We would also like to thank Tony Chen, Grain Legume Genetics Physiology Research, USDA ARS, Pullman, WA for assistance in laboratory work. We would also like to thank Derek Pouchnik and Weiwei Du of Laboratory for Biotechnology and Bioanalysis Washington State University, Pullman, WA for their assistance with plasmid and DNA sequencing. Financial support from the Indian Council for Agricultural Sciences (ICAR) is gratefully acknowledged.

## 8 Supplementary Material

The supplementary material for this article can be found online at:

