## Supplementary Material for "Identification of two novel polygalacturonase-inhibiting proteins (PGIPs) and their genomic reorganization in chickpea (*Cicer arietinum*)"

SUPPLEMENTARY FIGURE 1: Standard curves generated by serial dilution of cDNA for 18SrRNA, 25SrRNA, *Capgip1*, *Capgip3*, and *Capgip4*. 1 microgram of cDNA diluted ten-folds. CT values are indicated on Y-axis. Dilution $Log_{10}$ is indicated on X-axis.

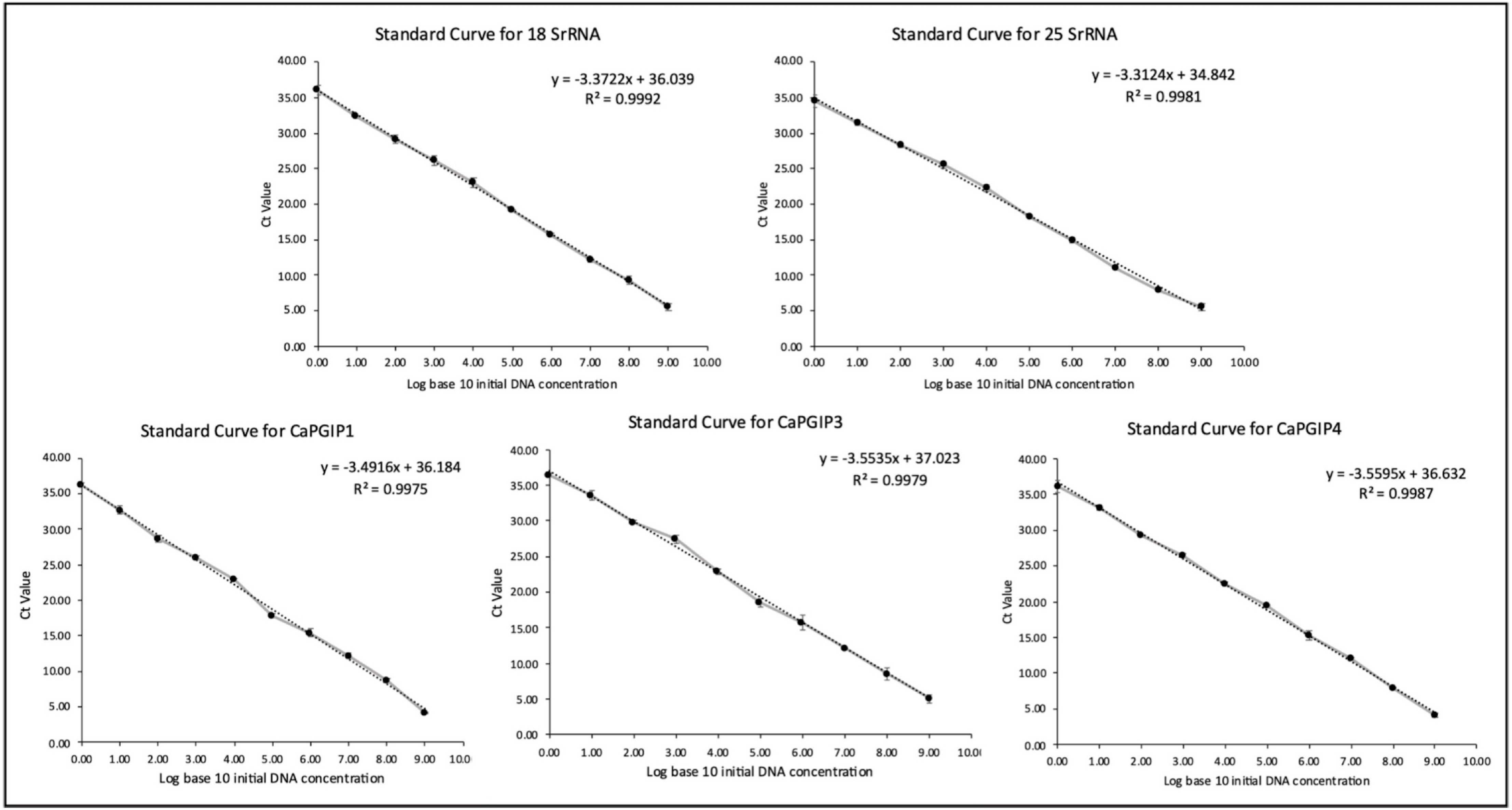

**SUPPLEMENTARY**

TABLE 1: Genes identified through a homology database search in *Cicer arietinum* genome that exhibited PGIP features.

| Sl. No. | Gene symbol | Location | Protein size (aa) | Signal peptide |
|---|---|---|---|---|
| 1 | LOC101488401 | Unplaced Scaffold | 138 | Absent |
| 2 | LOC101489504 | Unplaced Scaffold | 239 | Absent |
| 3 | LOC101499240 | Ca3 | 335 | Present |
| 4 | LOC101499557 | Ca3 | 329 | Present |
| 5 | LOC101504619 | Ca6 | 164 | Absent |
| 6 | LOC101505245 | Ca6 | 347 | Present |
| 7 | LOC105852278 | Ca6 | 225 | Absent |

TABLE 2: Primers used in this study to isolate, clone, and for the RT-qPCR expression analysis of the *Capgips*.

| **Gene** | **Primer name** | **Primer sequence (5' – 3')** | **Utility** |
|---|---|---|---|
| *Capgip1* | CaPGIP1-F | ATGAAGAACAAAATATTATCATCAT | Amplifying ORF |
| | CaPGIP1-R | AGTTTTGCAGGGTGGAAGAGGAGAA | |
| *Capgip3* | CaPGIP3-F | ATGGAAACAATATTAATAG | Amplifying ORF |
| | CaPGIP3-R | CTTGCAAGGCGGAAGTGGT | |
| *Capgip4* | CaPGIP4-F | ATGGCAACCGCTGTGCTAC | Amplifying ORF |
| | CaPGIP4-R | TGTACATTTGGGAAGCGGAGA | |
| *Capgip1* | CaPGIP1-q-F | GTAACCAACTCACCGGAGCA | RT-qPCR expression analysis |
| | CaPGIP1-q-R | GTCCAAAAACGCCAGGTTCA | |
| *Capgip3* | CaPGIP3-q-F | GATACCGCAGGGTGGTGAAT | RT-qPCR expression analysis |
| | CaPGIP3-q-R | TTCACTTGCAAGGCGGAAGT | |
| *Capgip4* | CaPGIP4-q-F | CAACTTCCCAACCTCAACGC | RT-qPCR expression analysis |
| | CaPGIP4-q-R | CCTAATGTGGGTGGGATGGG | |
| *18S ribosomal RNA* | 18SrRNA-F | ACGTCCCTGCCCTTTGTACAC | Reference gene for RT-qPCR expression analysis |
| | 18SrRNA-R | CACTTCACCGGACCATTCAAT | |
| *25S ribosomal RNA* | 25SrRNA-F | AAAACAAAGCATTGCGATGGT | Reference gene for RT-qPCR expression analysis |
| | 25SrRNA-R | GCACTGGGCAGAAATCACATT | |

TABLE 3: Other putative *cis*-acting regulatory elements identified in the promoter regions of *Capgips*.

| | ***Cis* - element** | **Position** | | | **Signal Sequence** | **Function** | **References** |
|---|---|---|---|---|---|---|---|
| | | ***Capgip1*** | ***Capgip3*** | ***Capgip4*** | | | |
| 1. | -10PEHVPSBD | - | 847 (+) | 171 (-) | TATTCT | Light response | Thum *et al.*, 2001 |
| 2. | ABREATRD22 | - | 284 (-) | - | RYACGTGGYR | Abscisic acid and dehydration response | Busk and Pagès, 1998; Iwasaki *et al.*, 1995 |
| 3. | CAATBOX1 | 15 (+),26 (+),48 (+),75 (+),102 (+),204 (-),294 (-), 338 (-),493 (-),512 (-),576 (+),839 (-), 908 (+),1011 (+), 1041 (-),1052 (+),1301 (-) | 77 (-), 123 (+), 150 (+), 217 (-), 223 (-), 245 (+), 293 (+), 298 (-), 451 (+), 481 (-), 496 (-), 650 (-), 673 (+), 688 (+),719 (-), 762 (-), 787 (+), 795 (-), 817 (-), 855 (+), 1061 (+), 1078 (+), 1235 (-), 1358 (-), 1374 (+),1463 (+),1475 (+) | 3 (+),64 (+),104 (+),137 (-),163 (+),230 (-),326 (+),415 (+),595 (-),622 (+),664 (-),681 (+),760 (-),817 (+),855 (-),948 (-),1081 (-),1207 (+),1228 (-),1328 (+),1383 (+),1461 (+) | CAAT | Common motif in promoter and enhancer regions responsible in tissue specific activity | Shirsat *et al.*, 1989 |
| 4. | CBFHV | 1212 (-) | 209 (+), 263 (+), 1160 (+) | 938 (-),938 (+),1087 | RYCGAC | DRE binding motif for dehydration response | Svensson *et al.*, 2006; Xue *et al.*, 2002 |
| 5. | CCAATBOX1 | 15 (+),26 (+),48 (+), 75 (+),102 (+),204 (-),294 (-),338 (-),493 (-),512 (-), 576 (+),839 (-),908 (+),1011 (+),1041 (-),1052 (+),1301 (-),1452 (-),1458 (+) | 1373 (+), 1462 (+) | 63 (+),162 (+),816 (+),1206 (+) | CCAAT | CAAT promoter motif commonly found in promoter and enhancer regions | Shirsat *et al.*, 1989 |
| 6. | CRTDREHVCBF2 | - | - | 938 (-),938 (+) | GTCGAC | Low temperature response | Xue 2003 |
| 7. | DPBFCOREDCDC3 | | - | 922 (+),1302 (+) | ACACNNG | DPBF-1 binding motif for abscisic acid response | Finkelstein *et al.*, 2000; Kim *et al.*, 1997; Lopez-Molina and Chua 2000 |
| 8. | DRE2COREZMRAB17 | 1212 (-) | 263 (+) | - | ACCGAC | Core site required for | Busk *et al.*, 1997; |

| | | | | | | | |
|---|---|---|---|---|---|---|---|
| | | | | | | binding of DRE proteins involved in ABA and drought response | Dubouzet *et al.*, 2003; Kizis and Pagès 2002; |
| 9. | DRECRTCOREAT | 1212 (-) | 263 (+) | 1087 (+) | RCCGAC | Core motif of DRE/CRT involved in drought, high-light, cold and heat stress response | Díaz-Martín *et al.*, 2005; Dubouzet *et al.*, 2003; Suzuki *et al.*, 2005; Qin *et al.*, 2004; |
| 10. | GATABOX | 9 (+),29 (-),86 (-), 236 (+), 245 (+), 356 (+), 380 (-), 392 (+),440 (+), 442 (-),470 (-),714 (+),843 (+), 1001 (-),1029 (+),1228 (+),1254 (+),1259 (+),1261 (-),1344 (-),1381 (+),1447 (-),1470 (-) | 56 (-), 159 (+), 166 (-), 198 (+), 205 (-), 220 (+), 257 (+), 429 (+), 522 (-), 670 (-), 712 (+), 835 (-), 870 (+), 1058 (-), 1070 (-), 1104 (-), 1118 (-), 1238 (+), 1447 (-), 1467 (-), 1472 (-) | 260 (+),409 (+),633 (+),974 (+),1048 (-),1128 (+),1145 (-),1264 (-),1322 (+),1390 (-),1458 (-) | GATA | Common cis-acting element in promoter for tissue specific expression | Gidoni *et al.*, 1989; Gilmartin *et al.*, 1990; Lam and Chua 1989; Reyes *et al.*, 2004; Rubio-Somoza at al., 2006; Teakle *et al.*, 2002; |
| 11. | LTREATLTI78 | 1211 (-) | 263 (+) | - | ACCGACA | Motif for low temperature response | Nordin *et al.*, 1993 |
| 12. | LTRECOREATCOR15 | 1212 (-) | 264 (+) | 1088 (+) | CCGAC | Core motif for low temperature response, drought induced gene expression ABA-regulated gene | Baker *et al.*, 1994; Jiang *et al.*, 1996; Busk and Pagès *et al.*, 1998; Kim *et al.*, 2002 |
| 13. | MYB1AT | 398 (-),762 (-),1048 (+) | 109 (+), 130 (+), 1492 (+) | 1012 (+) | WAACCA | MYB recognition site found in the promoters of the | Abe at al., 2003 |

| | | | | | | | |
|---|---|---|---|---|---|---|---|
| | | | | | | dehydration-responsive gene | |
| 14. | MYB2AT | 499 (+),699 (+),1235 (+) | - | 421 (+) | TAACTG | MYB recognition site for water stress response | Urao *et al.*, 1993 |
| 15. | MYB2CONSENSUSAT | 499 (+),595 (-), 699 (+),1235 (+) | - | 421 (+),1398 (-) | YAACKG | MYB recognition site for dehydration response | Abe *et al.*, 2003 |
| 16. | MYBCORE | 499 (-),595 (+), 699 (-),1214 (+), 1235 (-) | 1141 (-) | 421 (-),899 (-),1398 (+) | CNGTTR | MYB recognition site for water stress response | Lüscher and Eisenman 1990; Urao *et al.*, 1993; Solano *et al.*, 1990 |
| 17. | MYCATERD1 | - | 1296 (-) | 1303 (-) | CATGTG | MYC recognition site for dehydration, drought stress | Simpson *et al.*, 2003 |
| 18. | MYCATERD22 | - | 1296 (+) | 1303 (+) | CACATG | MYC recognition site for drought- and abscisic acid-regulated gene expression | Abe at al., 1997; Busk and Pagès *et al.*, 1998; |
| 19. | MYCCONSENSUSAT | 595 (-),595 (+), 1333 (-),1333 (+) | 287 (-), 287 (+), 467 (-), 467 (+), 707 (-), 707 (+), 1287 (-), 1287 (+), 1296 (-), 1296 (+), | 298 (-),298 (+),1123 (-),1123 (+),1303 (-),1303 (+),1398 (-),1398 (+),1417 (-),1417 (+) | CANNTG | MYC recognition site involved in dehydration and cold response | Abe *et al.*, 2003; Agarwal *et al.*, 2006; Chinnusamy *et al.*, 2003; Chinnusamy *et al.*, 2004; Hartmann *Et al.*, 2005; Lee *et al.*, 2005; Oh *et al.*, 2005; |

| | | | | | | | |
|---|---|---|---|---|---|---|---|
| 20. | QARBNEXTA | - | 1414 (-) | 560 (-),1131 (-),1166 (+) | AACGTGT | Motif for wounding and tensile stress response | Elliot and Shirsat, 1998 |
| 21. | OSE1ROOTNODULE | - | - | 971 (+),1146 (-) | AAAGAT | Activation in the infected cells of root nodules | Fehlberg *et al.*, 2006; Vieweg *et al.*, 2004; |
| 22. | OSE2ROOTNODULE | 279 (-),435 (+), 1308 (+) | - | - | CTCTT | consensus sequence motifs for promoter activation in the infected cells of root nodules | Fehlberg *et al.*, 2005; Vieweg *et al.*, 2004 |
| 23. | ROOTMOTIFTAPOX1 | 10 (+),27 (-),53 (-), 54 (+),156 (+),302 (-),332 (-),378 (-), 393 (+),428 (+), 658 (+),890 (-), 893 (+),928 (+), 944 (+),994 (+), 1030 (+),1119 (-), 1120 (+),1348 (+) | 4 (-), 9 (+), 51 (+), 115 (-), 215 (+), 221 (+), 344 (-), 345 (+), 404 (-), 435 (-), 440 (-), 441 (+), 518 (-), 587 (+),596 (-), 674 (-),689 (-),690 (+),753 (+),833 (-),845 (-),846 (+),1033 (+),1054 (-),1062 (-),1063 (+),1068 (-),1128 (-),1191 (-), 1192 (+), 1232 (-), 1233 (+), 1375 (-), 1376 (+), 1386 (+), 1404 (-), 1445 (-) | 11 (-),37 (-),40 (+),327 (-),328 (+),380 (+),440 (-),441 (+),469 (-),470 (+),475 (+),482 (-),634 (+),703 (-),704 (+),1042 (-),1043 (+),1388 (-), 1462 (-),1463 (+) | ATATT | Root specific expression | Elmayan and Tepfer., 1995 |
| 24. | RYREPEATBNNAPA | - | 79 (-) | - | CATGCA | Cis elements for ABA response | Ezcurra *et al.*, 1999; Ezcurra *et al.*, 2000 |
| 25. | TATABOX2 | 670 (+),766 (+), 793 (-),938 (-), 1032 (-) | 534 (+),1050 (+),1092 (+),1130 (+),1406 (+),1441 (+), | 306 (-),376 (-) | TATAAAT | Common *cis*-acting element responsible for the tissue specific promoter activity | Grace *et al.*, 2004; Shirsat *et al.*, 1989 |
| 26. | TATABOX3 | 834 (+) | 1064 (+),1065 (-), 1387 (+) | 142 (-),384 (+),1240 (-),1338 (+),1385 (-),1464 (+) | TATTAAT | Common *cis*-acting element in promoter and enhancer regions | Shirsat *et al.*, 1989 |

| | | | | | | | |
|---|---|---|---|---|---|---|---|
| 27. | TATABOX4 | 724 (+),940 (-),<br>1217 (-),1218 (+),<br>1247 (-) | - | 402 (-) | TATATAA | Common *cis*-acting element in promoter and enhancer regions | Grace *et al.*, 2004 |
| 28. | TATABOX5 | 93 (+),108 (+),120 (-),264 (-),285 (-),374 (-),419 (+),454 (+),673 (-),688 (+),791 (+),948 (+),<br>1115 (-),1181 (+),<br>1186 (+),1292 (+), | 17 (-),105 (-),543 (+),566 (+),635 (+),<br>639 (+),1095 (-),<br>1250 (-),1317 (+), | 287 (-),360 (+),364 (+),503 (-),685 (-),721 (-),755 (-),986 (+),1003 (-),1029 (+),1136 (+),1189 (-),1211 (-),1215 (-) | TTATTT | Common *cis*-acting element in promoter and enhancer regions | Tjaden *et al.*, 1995 |
| 29. | WBBOXPCWRKY1 | - | - | 1291 (+) | TTTGACY | WRKY binding site, involved in many plants physiological processes | Eulgem *et al.*, 2000 |
| 30. | WBOXATNPR1 | 197 (+) | 314 (-),484 (-),1043 (+) | - | TTGAC | WRKY binding site, involved in many plants physiological processes | Chen and Chen 2002; Chen *et al.*, 2002; Eulgem *et al.*, 2000; Maleck *et al.*, 2000; Yu *et al.*, 2001; |
| 31. | WRKY71OS | 198 (+), 1337 (+) | 314 (-),484 (-), 662 (-),809 (+),1044 (+) | 413 (-),781 (-),1061 (-),1119 (+),1293 (+),1300 (+),1307 (+),1370 (-) | TGAC | WRKY binding site, involved in many plants physiological processes | Eulgem *et al.*, 1999; Eulgem *et al.*, 2000; Xie *et al.*, 2005; Zhang *et al.*, 2004 |
